# Isolating syntax in natural language: MEG evidence for an early contribution of left posterior temporal cortex

**DOI:** 10.1101/439158

**Authors:** Graham Flick, Liina Pylkkänena

**Author notes:** Corresponding author: Graham Flick 10 Washington Place New York, NY 10003, USA.

## Abstract

Syntax is the engine that allows us to create an infinitude of linguistic expressions, and the construction of syntactic structures, such as noun phrases and verb phrases, is considered a fundamental component of language processing. Nevertheless, insights concerning the neurobiological basis of syntax have remained elusive, in part because it is difficult to isolate syntax from semantic composition. Consequently, many studies of syntax have relied on meaningless artificial stimuli, such as jabberwocky expressions or artificial grammars. However, while pure manipulations of syntax are challenging to design, natural language grammars do have a sparse set of constructions presenting this possibility. Here we examined one such case, English post-nominal adjectives (mountain TALL enough for a strenuous hike), which were contrasted with semantically parallel but structurally simpler noun-adjective sequences in an MEG experiment. We observed a sharp activity increase in the left posterior temporal lobe (PTL) when syntactic composition was more straightforward, approximately 200 ms after adjective onset. The semantic fit between the noun and adjective was also varied, but this affected anterior temporal cortex, consistent with prior work. These findings offer a unique demonstration of the relevance of posterior temporal cortex for syntactic processing in natural language. We also present connectivity evidence that the syntax-related PTL responses were relayed to ipsilateral inferior frontal and anterior temporal regions. The combined results draw an initial picture of the rapid spatio-temporal dynamics of the syntactic and semantic composition network in sentence processing.

## 1. INTRODUCTION

The construction of syntactic structures, such as noun phrases, verb phrases and, ultimately, sentences, is widely considered a fundamental component of language processing. What neurobiological mechanisms underlie this process? Answers to this question have remained elusive, in part because, theoretically, syntactic composition (e.g., the combination of an adjective and a noun to yield a noun phrase, identical in both *red fox* and *dead fox*) is difficult to isolate from processes related to semantic composition (i.e., the construction and combination of complex meanings, yielding quite different results in *red fox* and *dead fox*). In fact, whether the brain performs a purely structural, syntactic process remains an open question, and investigations into this matter have typically relied on inferences from combinatory processing more generally, or artificial linguistic stimuli. As such, there is a paucity of evidence concerning the neurobiological instantiation of syntactic composition, adequately isolated from its semantic counterpart, during the comprehension of natural language.

Much of the work that informs neurobiological accounts of syntax comes from more general manipulations of sentence or combinatory processing. As described by Friederici (2011), common approaches have included the contrast of sentence and word-list conditions, the introduction of syntactic violations, or various manipulations of syntactic complexity (e.g. object versus subject relative clauses). The results of such manipulations have pointed to several important left hemisphere regions, typically housed in the left anterior temporal lobe (ATL), the left posterior temporal lobe (PTL; as well as the neighboring angular gyrus and temporo-parietal junction), and the left inferior frontal gyrus, which may underpin syntactic aspects of language comprehension.

More recent work has corroborated the significance of these regions in relevant aspects of comprehension. Focusing on the left PTL, previous work on brain responses to syntactic and semantic ambiguities have suggested that sections of the PTL underlie the retrieval of lexico-syntactic information from memory (Snijders, Vosse, Kempen, Van Berkum, Petersson, & Hagoort, 2009; Rodd, Longe, Randall, & Tyler, 2010; Tyler, Cheung, Devereux, & Clarke, 2013), and/or a process of syntactic prediction employed during sentence processing (Matchin, Hammerly, & Lau, 2017; Matchin, Brodbeck, Hammerly, & Lau, 2018). Left PTL responses to naturalistic language stimuli (e.g., audiobooks) have also been shown to correlate with the predictions of hierarchical syntactic models (Brennan et al., 2016), and a recent investigation of agrammatical comprehension linked only the left PTL and nearby inferior parietal cortex with the processing of complex syntactic structure (Rogalsky et al., 2018).

The IFG, although not historically found in studies contrasting sentences and lists of words (see Friederici, 2011), has recently been shown to house increased activation in response to sentences if the lists contain only function or content words (Zacarella, Schell, & Friederici, 2017), and increases in response to individual instances of composition in adjective-pseudo-word phrases (e.g., *this FLIRK*, where *FLIRK* is a pseudo-word) relative to lists composed of noun-pseudo-word combinations (e.g., *apple, FLIRK*; Zaccarella & Friederici, 2015). While not finding effects in the IFG, a series of MEG studies have demonstrated that the left ATL shows increased activation in response to individual instances of composition in phrases, though this has predominantly used a consonant string + noun baseline condition (*red boat* vs. *xwk boat*), with composition-related increases appearing approximately 200 ms following the onset of the phrasal head (Bemis & Pylkkänen, 2011, 2012, 2013a, 2013b). Furthermore, like the PTL, ATL responses have been found to correlate with the predicted costs of integration from hierarchical syntactic models (Brennan, Nir, Hasson, Malach, Heeger, & Pylkkänen, 2010; Brennan et al., 2016; Brennan & Pylkkänen, 2017).

While valuable, the contribution of these findings to a neurobiological account of syntax is limited by how well the manipulations isolate syntax, per se. For instance, as is readily acknowledged, the sentence versus list contrast differs in both syntactic and semantic aspects of composition. Furthermore, the use and manipulation of syntactic complexity typically co-occurs with changes in sentence meaning, working memory demands (see Santi & Grodzinsky, 2007 for related discussion), or the order in which word meanings are recognized and combined (an issue for the delayed hemodynamic response, which blurs together a complete sentential response). The more recent trend of using naturalistic stimuli and models of syntactic parsing is also not without limitation, since increases in syntactic complexity may correlate with increases in, for example, conceptual specificity (Brennan et al., 2016). Those focusing on single instances of composition (e.g. Bemis & Pylkkänen, 2011; Zaccarella & Friederici, 2015) also face the challenge of teasing apart the semantic and syntactic aspects of the combinations, as illustrated by the *red fox* and *dead fox* example. Further to this point, the ATL composition response found in MEG studies of minimal composition appears to be more consistent with a semantic, rather than syntactic, composition process (Westerlund & Pylkkänen, 2014; Zhang & Pylkkänen, 2015; Poortman & Pylkkänen, 2016; Ziegler & Pylkkänen, 2016), arguing against the characterization of this region as a syntactic composition engine.

In response to these difficulties, others have attempted to subtract out lexical-semantic processing by turning to artificial or meaningless linguistic stimuli. One prominent example is studies that ask participants to learn artificial grammars specifying acceptable/unacceptable sequences of syllables. Such studies have associated sections of left inferior frontal, and ventral prefrontal, cortex with the processing of phrase structure rules (Opitz & Friederici, 2003, 2004) and, more specifically, hierarchical dependencies that are proposed to exist in syntactic structure (Friederici, Bahlmann, Heim, Schubotz, & Anwander, 2006; Bahlmann et al., 2008; see also Makuuchi, Bahlmann, Anwander & Friederici, 2009). Another example is a variant of the sentence versus word list paradigm, in which content words are replaced with pseudo-words, creating jabberwocky sentences (see Humphries, Binder, Medler, & Liebenthal, 2006 for example) and their scrambled counterparts (jabberwocky lists). The results of such studies have been mixed. Earlier work demonstrated increased activation in response to jabberwocky sentences relative to pseudo-word lists in the left ATL (Mazoyer et al., 1993; Friederici, Meyer, & von Cramon, 2000; Humphries et al., 2006). However, more recent investigations have failed to show the same result, instead implicating sections of the left PTL and inferior frontal cortex in syntactic composition (Pallier, Devauchelle, & Dehaene, 2011) or prediction (Matchin et al., 2017), or suggesting that there is no single region selectively sensitive to syntax (Fedorenko, Nieto-Castañon, & Kanwisher, 2012). Crucially however, the Jabberwocky sentences likely trigger a form of semantic composition that builds a coarse event depiction, allowing one to readily infer, for example, the roles of the agent, patient and action, even though these are occupied by pseudo-words. Correlates of said process are therefore expected to appear alongside any potential component related to pure syntactic composition, reducing the functional specificity of the conclusions regarding activation in the PTL and IFG. As regards artificial grammar learning, similar to violation studies, it is unclear whether the strategies adopted by individuals in this context generalize to natural comprehension.

In sum, approaches to the study of composition in the brain have pointed to sections of the left PTL and the left IFG as potentially housing computations critical to syntactic composition, with some results suggesting that each region, or smaller sub-regions, may underlie specific sub-processes (e.g., hierarchical dependency processing in the IFG). The ATL, on the other hand, has been implicated in semantic aspects of composition (Westerlund & Pylkkänen, 2014; Ziegler & Pylkkänen, 2016; Poortman & Pylkkänen, 2016; c.f. Baron & Osherson, 2011; Boylan, Trueswell, & Thompson-Schill, 2017), though a role in syntax has also been suggested (Brennan et al., 2010). However, the functional characterizations of these regions have been based predominantly on studies that either inferred the syntactic nature of correlates without adequately ruling out other computations, and/or relied on the use of artificial stimuli, which may trigger processing strategies distinct from those that support natural comprehension. While the inherent difficulty in manipulating syntactic composition independent of semantic composition makes this situation a reasonable one, the confidence with which any given correlate can be associated with syntactic computation, in particular, could be further enhanced through studies that manipulate syntax, independent of semantic composition, in natural language. Crucially, there are corners of natural language grammars in which pure manipulations of syntactic structure are at least arguably possible. In the present investigation, we took advantage of one such case in English, with one half of the contrast involving post-nominal modification.

In English, the canonical position of adjectival modifiers is pre-nominal (e.g., *There are many wide trails*), with post-nominal positioning leading to ungrammaticality in most cases (*There are many trails wide*). However, ‘heavy’ modifiers, such as those that appear with a complement (*There are many trails wide enough for a bear*), must appear post-nominally. This entails that, when encountered incrementally, the post-nominal adjective does not grammatically compose with its context until the complement is encountered. Crucially, we can design a contrasting instance of natural language that is almost identical, lexically, but differs in the syntactic composability of the adjective. To do so, we started with the same set of words, but with the adjective functioning as the main predicate as opposed to the modifier of the noun (*many trails are wide…*). To accomplish adjacency between noun and adjective, we employed an interrogative frame of the predicate structure (*are many trails wide enough…*), forming a four-word string-identical sequence with the post-nominal modification structure (*there are many trails wide enough…*). As will likely be noted right away, the processing of declarative and interrogative sentences confounded this contrast. While this is unavoidable, it was straightforward to build a control comparison that may account for the difference, by embedding the same combinations of adjectives and nouns, in their canonical order, in similar declarative and interrogative sentences that do not differ in the syntactic operation of interest.

Here, we adopted this comparison in an MEG study aimed at localizing, in both space and time, the correlates of the distinct syntactic combinatory possibilities of the critical adjectives. In addition, we simultaneously manipulated the match between the semantic types of the adjectives and nouns. This allowed us to demonstrate that the resulting correlate showed an insensitivity to semantic properties of the composing words, as would be expected of a purely syntactic component of processing. It also enabled us to search for separate correlates of semantic composition, which previous work has associated with the left anterior temporal lobe (e.g. Westerlund & Pylkkänen, 2014), and thus provide a spatiotemporal characterization of combinatorics at both levels of representation. Finally, we also performed connectivity analyses between our areas of interest, creating a link between the present results and accounts that posit an interaction between the left posterior temporal lobe and inferior frontal cortex that underpins syntactic composition (e.g., Den Ouden et al., 2012; Griffiths, Marslen-Wilson, Stamatakis, & Tyler, 2013; Schoffelen, Hultén, Lam, Marquand, Uddén, & Hagoort, 2017).

## 2. MATERIALS AND METHODS

### 2.1 Participants

This investigation was split across two separate experiments, conducted with a largely shared sample of participants, with data collected on separate days. Fourteen participants took part in each experiment (Experiment 1: 11 females, mean age = 25.58 years, standard deviation = 9.327 years; Experiment 2: 12 females, mean age = 21.83 years, sd. = 1.691). Twelve of those fourteen participants took part in both experiments, with two additional participants unique to each. The order of experiments was balanced across those participants that completed both. All participants were native English speakers, with healthy or corrected-to-healthy vision, and healthy hearing. All participants were right-handed.

### 2.2 Design

The structure of the Main experiment is displayed in Table 1. The materials consisted of sentences containing post-nominal modification (*There are many trails wide…*), and sentences containing string-identical sequences that, rather than modification, exemplified predication structures (*Many trails are wide…*) in an interrogative form (A*re many trails wide…*). A visual depiction of this contrast is shown in Figure 1. To reveal the semantic similarity between the pair of modification and predication, one can paraphrase the modification structure with the insertion of a relative clause (*There are many trails that are wide enough for a bear*), and compare this to the declarative form of the predication (*Many trails are wide enough for a bear*). This reveals that both sentences refer to some subset of all individuals (i.e. many of all existing trails) and pairs them with a given property (i.e. are of sufficient width). The remaining distinctions between the conditions used in the experiment can then be characterized as: (i) the syntactic integration that takes place on the noun-trailing adjective and (ii) the contrast between declarative and interrogative form. The former is the process that we intended to isolate, while the latter is a confounding difference that is, in this context, unavoidable.

**Figure 1.**
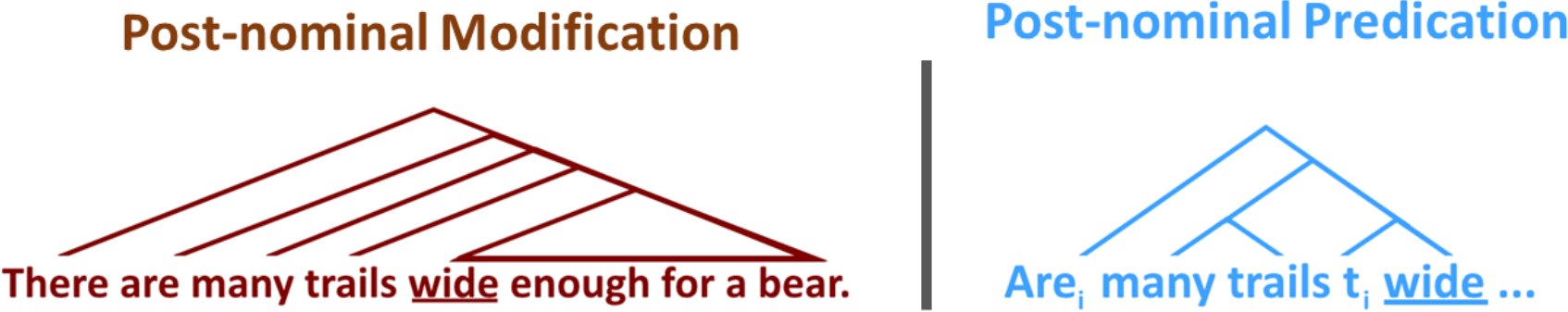
A tree visualization of the Syntactic Frame manipulation, showing the difference in syntactic composability. In post-nominal modification (left), the adjective requires the complement that follows it to grammatically compose with the preceding noun. In contrast, the predication relationship (right; depicted here with the use of a trace marker: t_i_) enables immediate and straightforward syntactic composition on the post-nominal adjective.

**Table 1.**
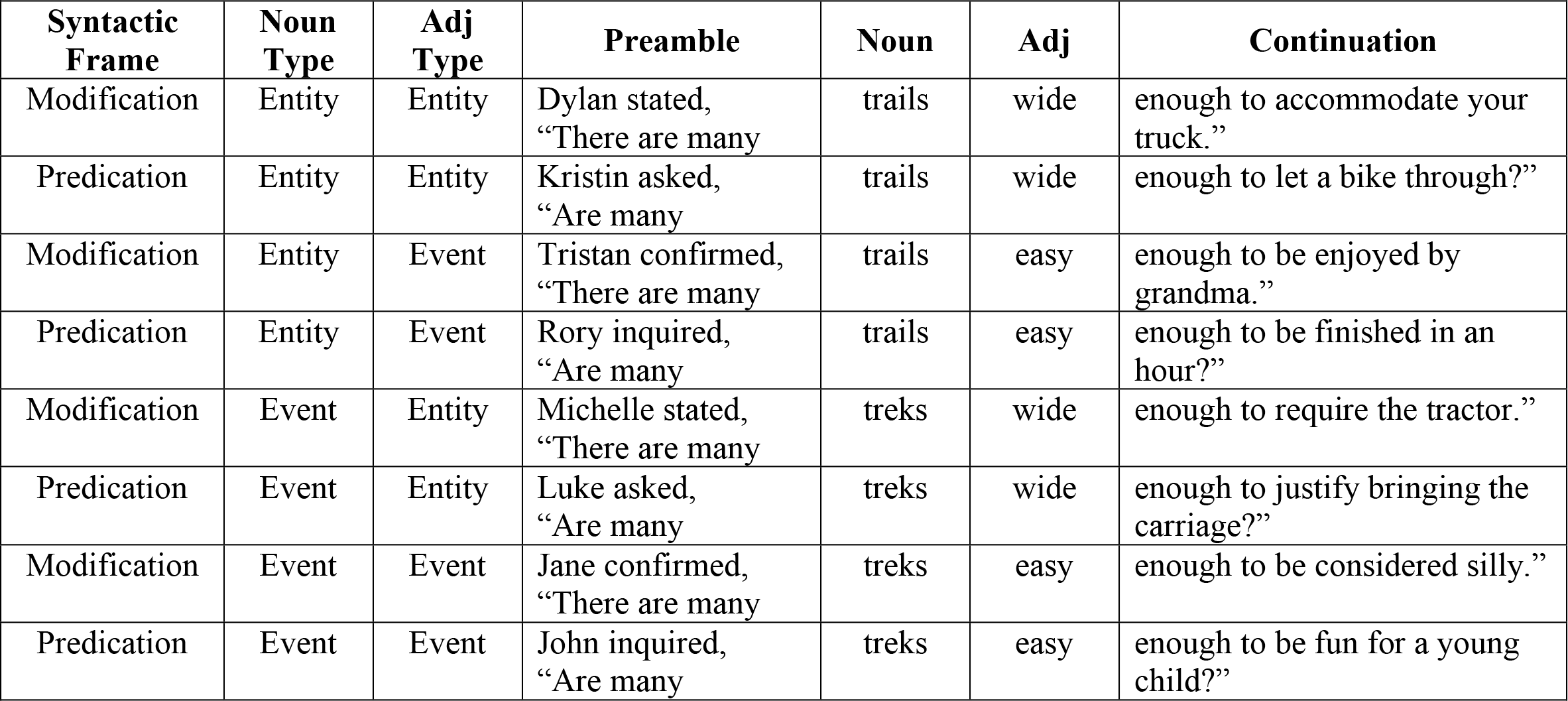
Half of one complete set of sentential stimuli, demonstrating a full crossing of Syntactic Frame, Noun Type, and Adjective Type in the Main Experiment.

As a control for this confound, we also tested for a general sensitivity to declarative versus interrogative sentences, using maximally similar lexical material. To do so, we embedded the same combinations of adjectives and nouns in secondary declarative (*There are many wide tails in…*) and interrogative (*Are there many wide trails in…*) sentences, this time using the canonical adjective-noun word order. This set of sentences, exemplified in Table 2, comprised the stimulus set in the second experiment, referred to as the Control Experiment.

**Table 2.** The second half of the set displayed in Table 1, containing the corresponding sentences used in the Control Experiment.

| **Sentence Type** | **Noun Type** | **Adj Type** | **Preamble** | **Adj** | **Noun** | **Continuation** |
| --- | --- | --- | --- | --- | --- | --- |
| Declarative | Entity | Entity | Christine stated,<br>“There are many | wide | trails | in the parks of Canada.” |
| Interrogative | Entity | Entity | Anthony asked,<br>“Are there many | wide | trails | in your cabin’s neighborhood? |
| Declarative | Entity | Event | Mary confirmed,<br>“There are many | easy | trails | in the forest ahead.” |
| Interrogative | Entity | Event | Allen inquired,<br>“Are there many | easy | trails | in the highlands near town?” |
| Declarative | Event | Entity | Sarah stated,<br>“There are many | wide | treks | in the area around Tuscon.” |
| Interrogative | Event | Entity | Gary asked,<br>“Are there many | wide | treks | in the surrounding valley? |
| Declarative | Event | Event | Joan confirmed,<br>“There are many | easy | treks | in the book on the shelf.” |
| Interrogative | Event | Event | Lucy inquired,<br>“Are there many | easy | treks | in the plan for next week?” |

We also varied the match of the semantic types called for and denoted by adjectives and nouns, respectively. Noun stimuli were classified as denoting entities (e.g. *trail*, *mountain)* or events (e.g. *trek*, *exercise*). Similarly, adjectives were classified as those that select for events (*easy*, *difficult*) and those that can straightforwardly modify entities (e.g. *wide*, *massive*). A previous eye-tracking study has suggested that a mismatch between the types of adjectives and nouns (e.g., *difficult mountain*, which is interpreted with the assertion of an event: a mountain that is difficult to climb), elicits online processing costs (Frisson, Pickering, & McElree, 2011). Such instances of enriched composition (see Pustejovsky, 1991 for related discussion) are considered similar to cases of complement coercion (Traxler, Pickering, & McElree, 2002), which have been characterized as a semantic composition process (Pylkkänen & McElree, 2006). In our design, we adopted an extension of the adjective-noun pairing manipulation employed by Frisson et al., fully crossing the factors of Noun Type (Entity, Event) and Adjective Type (Entity, Event). The complete set of adjective-noun stimuli can be found in Appendix A. Appendix B contains the descriptions and results of 2 norming studies that confirmed the entity/event distinction between the noun stimuli, and the coercive nature of the event adjective, entity noun pairings.

In summary, each of the two experiments presented here consisted of a 2 × 2 × 2 factorial design. In the main experiment, which contained all sentences with noun-adjective word order, these factors were Noun Type (Entity, Event), Adjective Type (Entity, Event), and Syntactic Frame (Modification, Predication). In the Control Experiment, Noun Type and Adjective Type were again manipulated, but the more general factor of Sentence Type (Interrogative, Declarative) replaced Syntactic Frame (see Tables 1 and 2).

### 2.3 Stimuli

We selected 39 entity-denoting nouns, each of which was paired with an entity adjective that modified it in a straightforward manner, without the need for semantic coercion (e.g. *wide trail*). We selected a set of 12 such adjectives and paired each with three or four entity nouns. We then selected a second set of 12 adjectives that, when combined with the entity nouns, required coercion of the entity noun to an eventive interpretation (e.g. *easy trail*), with each adjective once again paired with three, or four entity nouns. Finally, to fully cross the factors, we generated a second set of 39 nouns denoting events, and matched each one to the appearance of an entity noun, therefore pairing with both of its corresponding adjectives. This resulted in 39 sets of adjective-noun quadruples. Notably, this complete crossing yields pairs of event nouns and entity adjectives that may be anomalous (e.g. *wide trek*) or, in some cases, involve a different sense of the adjective (e.g. *massive test*). Since there was a considerable degree of heterogeneity within this cell of the design, and the MEG results showed no modulation that could be straightforwardly associated with this cell in particular, we do not discuss these materials in any further detail.

The final sets of nouns were selected such that independent t-tests between them failed to show a statistically significant difference on the following measures: the logarithmic transform of the Hyperspace-to-Analog (HAL) frequency value, average letter bigram frequency, and the number of letters, phonemes, syllables, and morphemes (all p-values > 0.20). This was also true of the two sets of adjectives (all p-values > 0.35). Additionally, the final stimulus set was designed such that the three conditions of adjective-noun combinations that were not potentially anomalous (i.e. *wide trail, easy trail, easy trek*, but not *wide trek*) did not differ on transition probability in either word order (p-values > 0.15). The two sets of entity and event adjectives were also balanced on their transition probabilities to the word ‘enough’ (p > 0.50) since this followed them in every sentence of the Main Experiment. Lexical frequency values were extracted from the CELEX corpus (Baayen, Piepenbrock, & Gulikers, 1995), while HAL frequency values and other lexical characteristics were extracted from the English Lexicon Project (Balota et al., 2007). Transition probability was calculated by dividing the bigram frequency of each pair of words by the unigram frequency of the initial word, in each word order. Frequencies were extracted from the Corpus of Contemporary American English (Davies, 2008) and transition probability was calculated following add-one smoothing (see Martin & Jurafsky, 2009). Lexical and phrasal characteristics are displayed in Tables 3 and 4.

**Table 3.** Lexical characteristics of nouns and adjectives. Mean values for each measure are reported, with standard deviation in parentheses. HAL = Hyperspace-to-analogue proxy of lexical frequency.

| Class | Type | HAL | Bigram | Length | Phoneme | Syllable | Morpheme |
| --- | --- | --- | --- | --- | --- | --- | --- |
| Noun | Entity | 9.07<br>(1.54) | 3363<br>(1727) | 5.79<br>(1.75) | 4.67<br>(1.75) | 1.85<br>(0.74) | 1.33<br>(0.58) |
|  | Event | 9.46<br>(1.84) | 3902<br>(2104) | 6.24<br>(1.84) | 5.16<br>(1.60) | 2.03<br>(0.88) | 1.50<br>(0.65) |
| Adj | Entity | 9.24<br>(1.88) | 3408<br>(1294) | 6.17<br>(1.40) | 4.83<br>(1.11) | 1.75<br>(0.75) | 1.50<br>(0.52) |
|  | Event | 9.54<br>(1.52) | 3528<br>(1417) | 6.83<br>(2.33) | 5.33<br>(1.67) | 2.00<br>(0.74) | 1.67<br>(0.49) |

**Table 4.** Transition probabilities (TP) between adjectives and nouns, in both orders, for the three conditions of interest. Mean values for each measure are reported, with standard deviations in parentheses.

| Type Pairing:<br>Adj, Noun | Transition Probability:<br>Adj to Noun | Transition Probability:<br>Noun to Adj |
| --- | --- | --- |
| Entity, Entity | $1.27 \times 10^{-6}$ ($1.43 \times 10^{-6}$) | $3.88 \times 10^{-7}$ ($2.25 \times 10^{-7}$) |
| Event, Event | $1.24 \times 10^{-6}$ ($1.81 \times 10^{-6}$) | $6.34 \times 10^{-7}$ ($1.19 \times 10^{-6}$) |
| Event, Entity | $8.93 \times 10^{-7}$ ($1.13 \times 10^{-6}$) | $4.13 \times 10^{-7}$ ($2.09 \times 10^{-7}$) |
| Entity, Event | -- | -- |

Finally, each of the pairings of adjective and noun were embedded in the two levels of Syntactic Frame and Sentence Type. For each participant, each quadruple of adjectives and nouns was paired with one of two quantifiers (*many* or *some*) that preceded it in the sentences. The number of times a certain quantifier was paired with a given set, and thus a certain cell of the design, was balanced across participants. Sentential material following the pair of critical words was variable to reduce any potential cueing effects (see Continuation in Tables 1 and 2), with two exceptions. First, the word immediately following the post-nominal adjectives in the Main Experiment was always ‘enough’. Second, the word immediately following the nouns in the Control Experiment was one of three prepositions (*on, in, along*), and this was held constant within individual sets of adjectives and nouns.

To move the critical adjective-noun pairs further from the onset of the sentences, and therefore reduce potential effects related to the depth of the material, the sentences were embedded as the object of two speech production verbs. These verbs matched the declarative (*stated, confirmed*) or interrogative (*asked, inquired*) form of the sentences. For each participant, a unique pairing of each speech verb to each of the 39 sets of adjective-noun quadruples was generated, with an approximately uniform distribution of pairings across the cells of the design (i.e. ‘inquired’ was paired with each level of Adjective Type and Noun type approximately the same number of times). Speech verbs were preceded by one of sixteen unique names that were randomly assigned to the sentences, for each participant.

### 2.4 Procedure

Prior to beginning the MEG recording, each participant’s head shape was digitized using a Polhemus FastSCAN system (Polhemus, Vermont, USA). Participants were instructed that their task was to read the sentences carefully as they appeared word-by-word on the screen. They were told that these sentences were designed to be examples of natural language, which they may encounter in a story or narrative, and that following random trials they would be asked a question about the immediately preceding sentence. The instructions indicated that this question could ask about something simple, such as whether a specific word was present, or something more subjective, such as the greater context in which the speech act may have occurred, and that there may not be an objectively correct answer. Each comprehension question was followed by the presentation of two possible answers, presented on either side of the screen, and participants responded by pressing one of two buttons with their left hand.

Participants completed the experiment lying supine in a dimly lit magnetically shielded room (MSR). Sentences were back-projected onto the center of a screen approximately 80 cm away from the participant. Each word was presented in black, size 12, Times New Roman font on a grey background, and subtended a vertical visual angle of 0.64 degrees. Each trial began with the appearance of a fixation cross for 300 ms, followed by an inter-stimulus interval (ISI) of 300 ms. The first word in every trial was the proper name beginning the stimulus item, presented for 300 ms, which was followed by a blank screen for a 300 ms ISI. Next, the speech verb was presented for 300 ms, with a 500 ms ISI following it to mimic the tendency to pause before speech content, as indicated by the presence of a comma in written language (i.e. Luke said, ‘There are…’). All subsequent words were presented for 300 ms with 300 ms ISIs between them.

Continuous MEG data were recorded at a sampling rate of 1000 Hz with a 208-channel axial gradiometer whole-head system (Kanazawa Institute of Technology, Kanazawa, Japan). During acquisition, high-and low-pass filters were applied at 0.1 and 200 Hz, respectively. The positions of five electromagnetic coils, attached to the forehead and next to each tragus, were measured relative to the location of the MEG sensors, once before and once following the experiment. Participants required approximately 70 minutes to complete each experiment.

### 2.5 MEG Data Preprocessing

MEG data were first noise-reduced using the Continuously Adjusted Least-Squares Method (Adachi, Shimogawara, Higuchi, Haruta, & Ochiai, 2001) available in the MEG 160 software (Yokogawa Electrical Corporation and Eagle Technology Corporation, Japan). All remaining data preprocessing steps and subsequent analyses were performed with the MNE-python (v.0.14; Gramfort et al., 2013, 2014) and Eelbrain (v. 0.25.3; DOI: http://doi.org/10.5281/zenodo.438193) python packages. The continuous data were low-pass filtered at 40 Hz. Independent Component Analysis was applied to each MEG recording to isolate and remove components that matched the profile of known artifacts (eye blinks, movement-related activity, and well-characterized external noise sources), using visual inspection for removal. The reconstructed MEG data were then split into epochs spanning 100 ms before the onset of the first word in the adjective-noun pair, to 600 ms following the onset of the second word (for a total duration of 1300 ms). Epochs containing amplitudes greater than an absolute threshold of 2000 fT were removed before further preprocessing. The remaining epochs in each of the 16 conditions were then down-sampled by a factor of 5 to make the following steps computationally tractable, and averaged to yield an evoked response for each condition, for each participant. For visualization of the extended time series in response to later words, the same parameters, and methods that follow, were used to generate the source estimates.

Cortically constrained estimates of source-level activity were computed from each participant’s evoked responses using dynamical statistical parameter mapping (dSPM: Dale et al., 2000). For each participant, and for each experimental session, the digitized head shape and fiducial landmarks were used to scale and co-register the “fsaverage” brain (available in the FreeSurfer software suite: https://surfer.nmr.mgh.harvard.edu) to the participant. A source space of 2562 sources per hemisphere was generated on the scaled cortical surface, and the boundary element model was employed to compute a forward solution. Channel noise covariance matrices were estimated from 150 segments of data, each 100 ms long, extracted from just before the onset of the fixation cross on each trial. To best ensure that these segments were artifact free, and centered at zero mean, we selected those 150 segments that had mean activity, across time and channels, closest to zero. Inspection of the standard deviation of the signal across all such pre-stimulus intervals confirmed that our selections were within a single standard deviation from zero, for every MEG recording. Covariance was estimated with an automated regularization method (see Engemann & Gramfort, 2015) to account for the relatively small number of samples. Inverse solutions were computed with “free” orientation and applied to each participant’s evoked responses to yield the estimates of source-level activity.

The primary analyses of MEG source estimates were performed on activity localized to four regions of interest (ROIs), displayed in Figure 2. These ROIs were defined from a previous study of constituent structure building (Pallier et al., 2011), which implicated several specific cortical regions, consistent with a large body of literature (see Hagoort & Indefrey, 2014; Friederici, 2011). We generated an initial set of regions by using the coordinates of the left temporal pole, anterior superior temporal sulcus, posterior superior temporal sulcus, temporo-parietal junction, and the pars orbitalis and pars triangularis, reported by Pallier et al. Each region was a sphere of 30 mm diameter around the reported point in Montreal Neurological Institute (MNI) space. This resulted in 6 initial regions. Due to the proximity of pairs of regions to each other, the relative spatial resolution of MEG compared to fMRI, and the issue of multiple comparison correction that comes with many ROIs, we combined pairs of regions that were contiguous in space. This yielded three of our four regions of interest, which we refer to as the Anterior Temporal Lobe (ATL), the Posterior Temporal Lobe (PTL), and the Inferior Frontal Gyrus (IFG). The fourth region was the left orbito-frontal cortex (ORB), which was included on the basis of its previous sensitivity to the processing of semantic type coercion (Pylkkänen & McElree, 2007). This region was defined as a combination of the left lateral and medial orbitofrontal labels from the “aparc” atlas (Desikan et al., 2006), available with Freesurfer.

**Figure 2.**
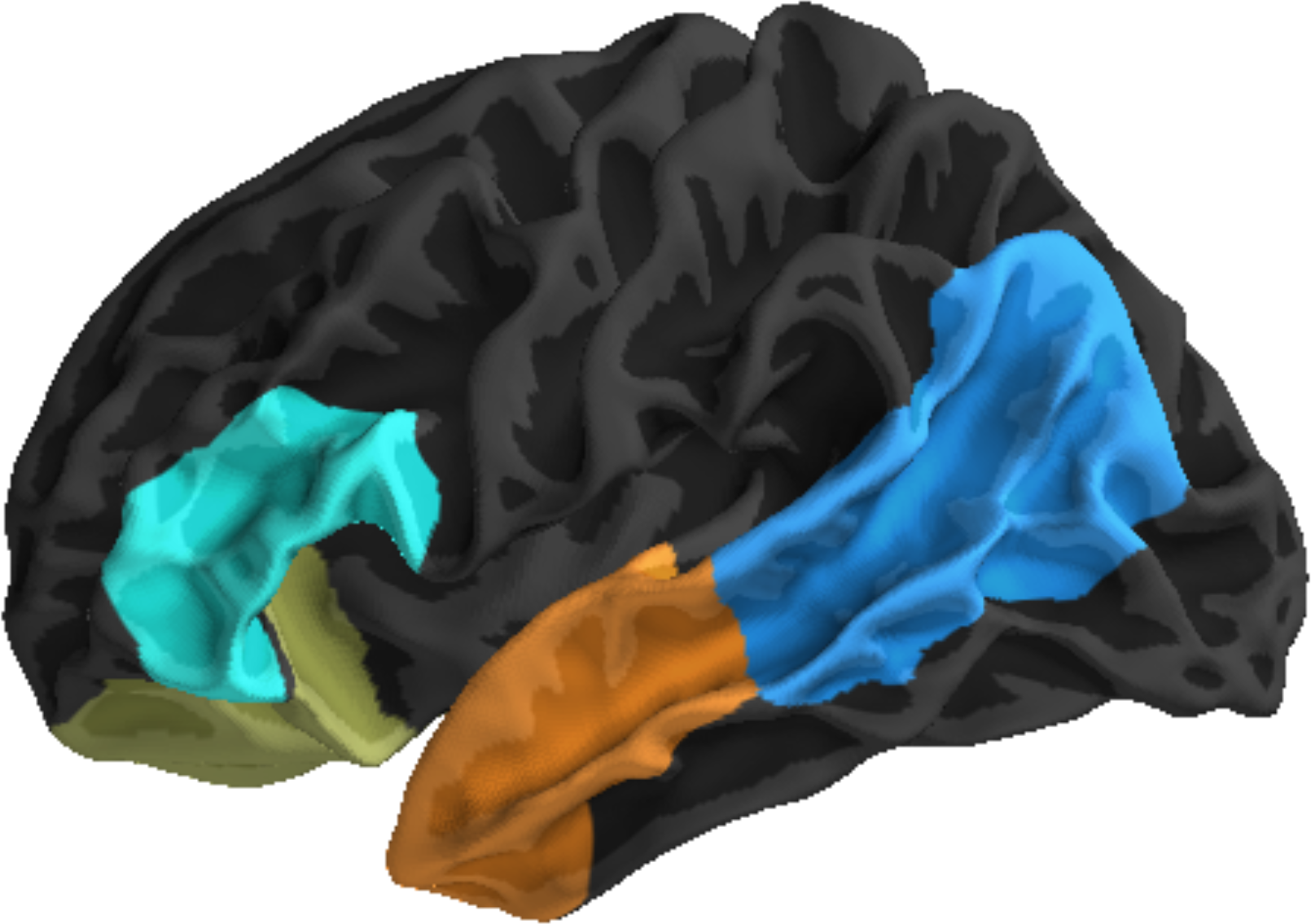
The anatomical regions of interest (ROIs) used in the factorial analysis of MEG source estimates.

We note that the combination of contiguous labels from Pallier et al. (2011) resulted in the pairing of two regions that the previous study characterized as sensitive, and insensitive, to the manipulation of lexical content: the temporo-parietal junction and posterior superior temporal sulcus, respectively. While this could reduce the present study’s sensitivity to related effects within each sub-region, we concluded that the reduction in the number of ROIs was more pressing. Furthermore, for each effect identified in the analyses that follow, we also performed a spatial assessment of its locus and report those in MNI coordinates, therefore allowing interested readers to compare the locations of the observed effects and use the positions in future research.

### 2.6 MEG Data Analysis

#### 2.6.1 Factorial Analysis

We first conducted cluster-based permutation tests (see Maris & Oostenveld, 2007 for method details and Bemis & Pylkkänen, 2011, for a similar usage) of activity localized to our four ROIs, in each experiment separately. The activity in each ROI was averaged over the sources at each time point, and the resulting time courses used as input in the temporal cluster-based tests. The statistic at each point in cluster formation was the F-value for the main effects and interactions in a 2 × 2 × 2 repeated measures analysis of variance (ANOVA) that included the factors Syntactic Frame (modification, predication), Noun Type (entity, event), and Adjective Type (entity, event). In the Control Experiment, Sentence Type (declarative, interrogative) replaced Syntactic Frame. Clusters were formed from statistics corresponding to a p-value less than 0.05, and only those clusters spanning a minimum of 20 ms were considered in the analysis. A permutation method using 10000 permutations was used to assign cluster p-values, which were corrected for multiple comparisons across the ROIs using the False Discovery Rate (Benjamini & Hochberg, 1995) with a critical value of 0.05. Our primary window of interest was on the responses to the second of the two critical words: the noun-trailing adjectives (trails *wide*) in the Main Experiment, and the noun (wide *trails*) in the Control Experiment. However, for completeness we also analyzed responses to the first word. The time window of analysis was 150-600 ms following the onset of each word.

For each temporal cluster that was identified in ROI time-courses, we used a secondary spatiotemporal cluster test to identify its spatial locus within the ROI. This was done by running the same cluster-forming procedure within the ROI over space as well as time, then identifying those spatiotemporal clusters that showed the same main effect or interaction within the time boundaries of the original cluster. The centers of these spatial clusters are reported in MNI coordinates, following a weighting of each source’s position by its average statistic across the duration of the spatiotemporal cluster.

#### 2.6.2 Connectivity Analysis

In addition to the factorial analyses of ROI activity, we also performed multivariate Granger Causality analysis (Granger, 1969; Barnett & Seth, 2014) on data collected in the Main Experiment. Granger Causality analysis can be understood as a model comparison approach to assessing whether activity in one region improves the prediction of activity in another target region, relative to prediction based on the target ROI’s own time course and those of all other ROIs included in the analysis. Significant degrees of improvement in prediction are taken to suggest a directional connection between two regions. We chose to use Granger Causality rather than other methods of estimating effective connectivity from MEG data (particularly, dynamic causal modeling; David, Kiebel, Harrison, Mattout, Kilner, & Friston, 2006), as it allowed us to analyze the same source estimates that were input to the factorial analysis. Based on the results of the factorial tests (see Results), we focused our primary assessment of Granger Causality (GC) on the predication condition of the Main Experiment. Nevertheless, we also present assessment of GC values in responses to adjectives in post-nominal modification for qualitative comparison.

For each participant, data in response to the adjectives (0-600 ms post word-onset) from the four ROIs was extracted and averaged across the pairings of Noun Type and Adjective Type. We also extracted evoked responses from left occipital cortex (defined as the lateral occipital label in the anatomical atlas of Desikan et al., 2006) to provide a more comprehensive model of responses to visual stimuli. The analysis was conducted using the Multivariate Granger Causality Matlab toolbox (Barnett & Seth, 2014), alongside all assumption checks and pre-processing steps described by Barnett and Seth (2014). For both Predication and Modification condition analyses, optimal model order was 5, from a possible space of orders between 1 and 20. In down-sampled time, this indicates a model that includes 25 ms prior, in each time series. Vector autoregressive (VAR) model parameters were estimated using ordinary least squares regression, with automatic determination of the maximum number of auto-covariance lags. Pairwise conditional Granger Causalities between regions were determined via log-likelihood comparisons of reduced and full models, wherein reduced models included all potential connections except the one being assessed. P-values for the resulting comparisons were determined via reference to the *F*-statistic distribution and corrected for multiplicity using the FDR procedure with a critical value of 0.05.

## 3. RESULTS

### 3.1 Main Experiment

Temporal, cluster-based 2 × 2 × 2 ANOVAs were performed on data localized to each of the four ROIs in response to the second of the two critical words (the adjectives). Two significant clusters were identified. A significant effect of Syntactic Frame, shown in Figure 3 (top-left) was found in the posterior temporal lobe (PTL) ROI, spanning 205-225 ms post-adjective onset (p = 0.0152). Inspection of the condition means revealed that the cluster contained increased activation in response to adjectives in post-nominal predication (mean = 1.251, sd = 0.299), relative to adjectives in post-nominal modification (mean = 1.143, sd = 0.269). Inspection of condition means across the levels of Noun Type and Adjective Type revealed that this pattern was present in every combination (Modification minus Predication: Entity, Entity: -0.1619; Entity, Event: -0.1602; Event, Entity: -0.1168; Event, Event: -0.2054).

**Figure 3.**
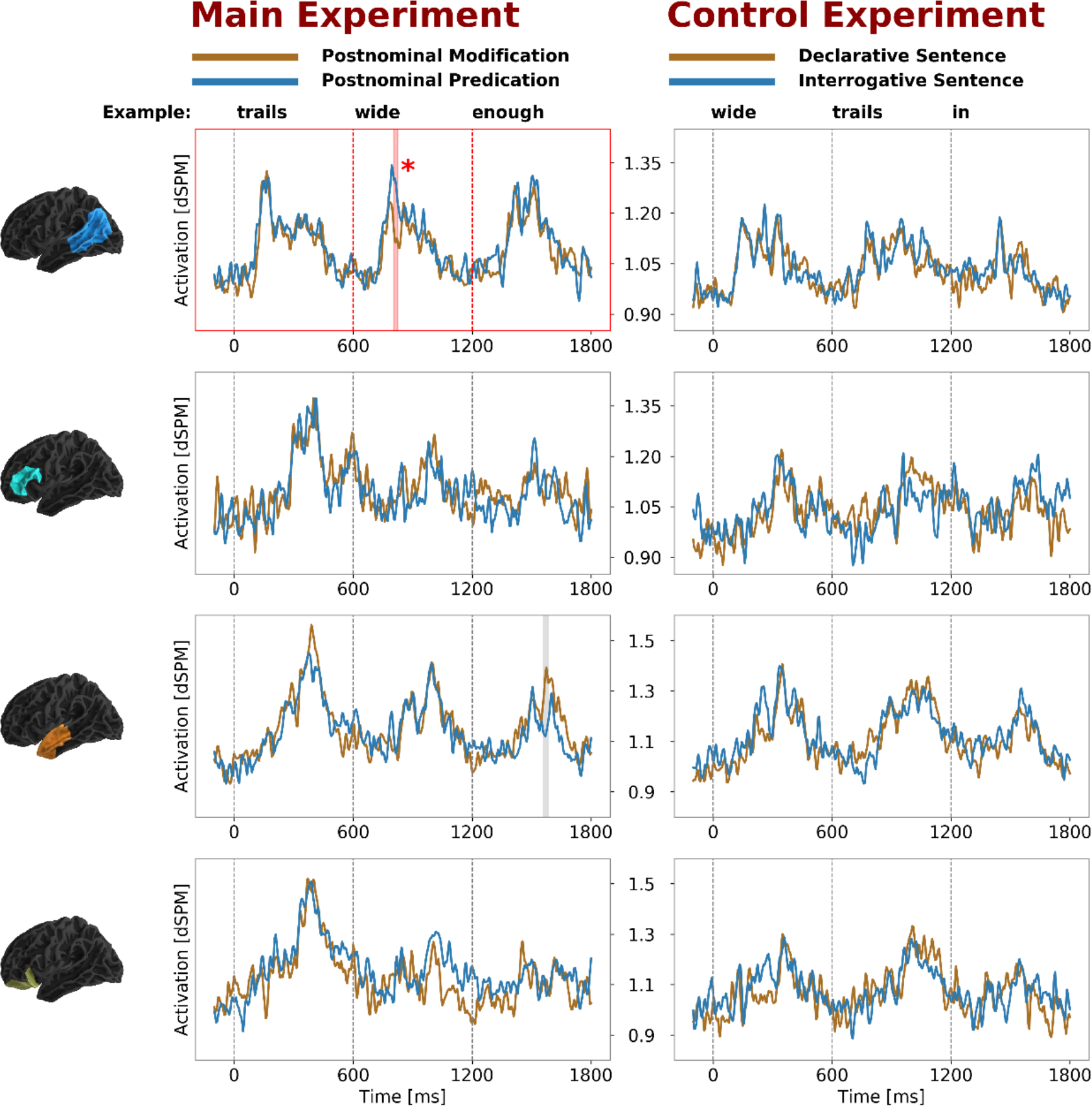
Main Experiment (left): A significant effect of Syntactic Frame was identified in the posterior temporal lobe, between 205 and 225 ms post-adjective onset. The cluster captured increased activation in instances of post-nominal predication, relative to post-nominal modification. No similar effect was identified in the Control Experiment.

A second significant cluster, displayed in Figure 4, was found in the ATL. The cluster spanned 175-195 ms (p = 0.0086) and captured a main effect of Noun Type, indicating an influence of the type of noun that preceded the adjective. Condition means demonstrated that the activation in the cluster was increased when adjectives followed an entity-denoting noun (mean = 1.215, sd = 0.388) relative to when they followed an event-denoting noun (mean = 1.015, sd = 0.231). Comparison of condition means across the levels of Syntactic Frame and Adjective Type revealed that this pattern was consistent across these factors, though responses were greatest, regardless of Syntactic Frame, when both noun and adjective were in the entity condition (Figure 4, bottom). No significant clusters (p > 0.05) were identified in response to the first of the two words, or the word ‘enough’ (which followed the noun), following correction for multiple comparisons.

**Figure 4.**
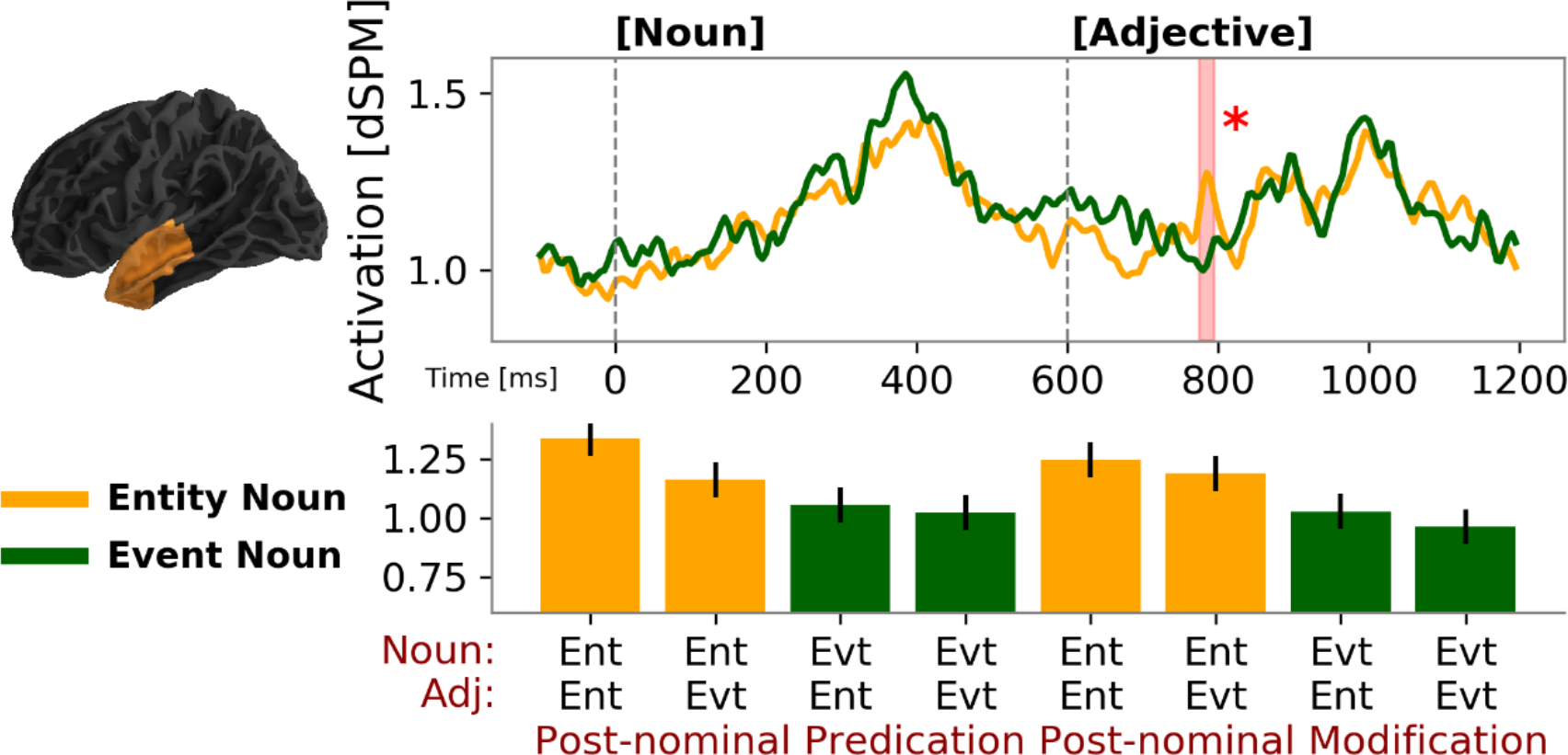
A main effect of Semantic Type of the noun was identified in responses to the following adjective in the Main Experiment. This appeared 175-195 ms post-adjective onset, in activity localized to the left anterior temporal lobe.

### 3.2 Control Experiment

A series of 2 × 2 × 2 ANOVAs were once again conducted to analyze responses to the first (adjective) and second (noun) critical stimuli in the Control Experiment. No significant clusters (p > 0.05) were found in response to either the adjective or the noun following correction for multiple comparisons. Crucially, this included a failure to find any semblance of the Syntactic Frame effect identified in the Main Experiment. To make this contrast clear, Figure 3 displays the parallel comparison of responses to declarative and interrogative sentences in every ROI. At no point in the time courses of any region were there significant F-values corresponding to the main effect of Sentence Type that met the cluster formation criterion of 20 ms duration. Exploratory analyses of responses to the prepositions (following the noun) also failed to find any statistically significant effects (p > 0.05).

### 3.3 Combined Factorial Analysis

Finally, we combined the data from the twelve participants who completed both the Main and Control Experiments in one dataset and analyzed this with a series of 2 × 2 × 2 × 2 repeated measures ANOVAs. For convenience, we recoded the factors as Word Order (Adjective-Noun, Noun-Adjective), Word 1 Type (Entity, Event), Word 2 Type (Entity, Event), and Sentence Type (Declarative, Interrogative). Two significant clusters containing main effects of Word Order were identified. The earlier of these two, displayed in Figure 5 (top) was found in response to the first critical word, in activity localized to the ORB ROI, 365-535 ms post word-onset (p = 0.0004). Inspection of condition means showed an increase in response to plural nouns (mean = 1.376, sd = 0.523) relative to adjectives in the same position (mean = 1.137, sd = 0.229). The second significant cluster was found in responses to the second critical word, in activity localized to the ATL, 450-520 ms post word-onset (p = 0.0044; Figure 5, bottom). Condition means again revealed an increase in response to nouns (mean = 1.320, sd = 0.388) relative to adjectives (mean = 1.150, sd = 0.269). The spatiotemporal loci of all significant effects were summarized by conducting the spatiotemporal cluster-based tests within each ROI, within the temporal boundaries of the clusters, and identifying those that matched the main effects of interest. These are displayed in Figure 6, along with the MNI coordinates of their mean-weighted centers.

**Figure 5.**
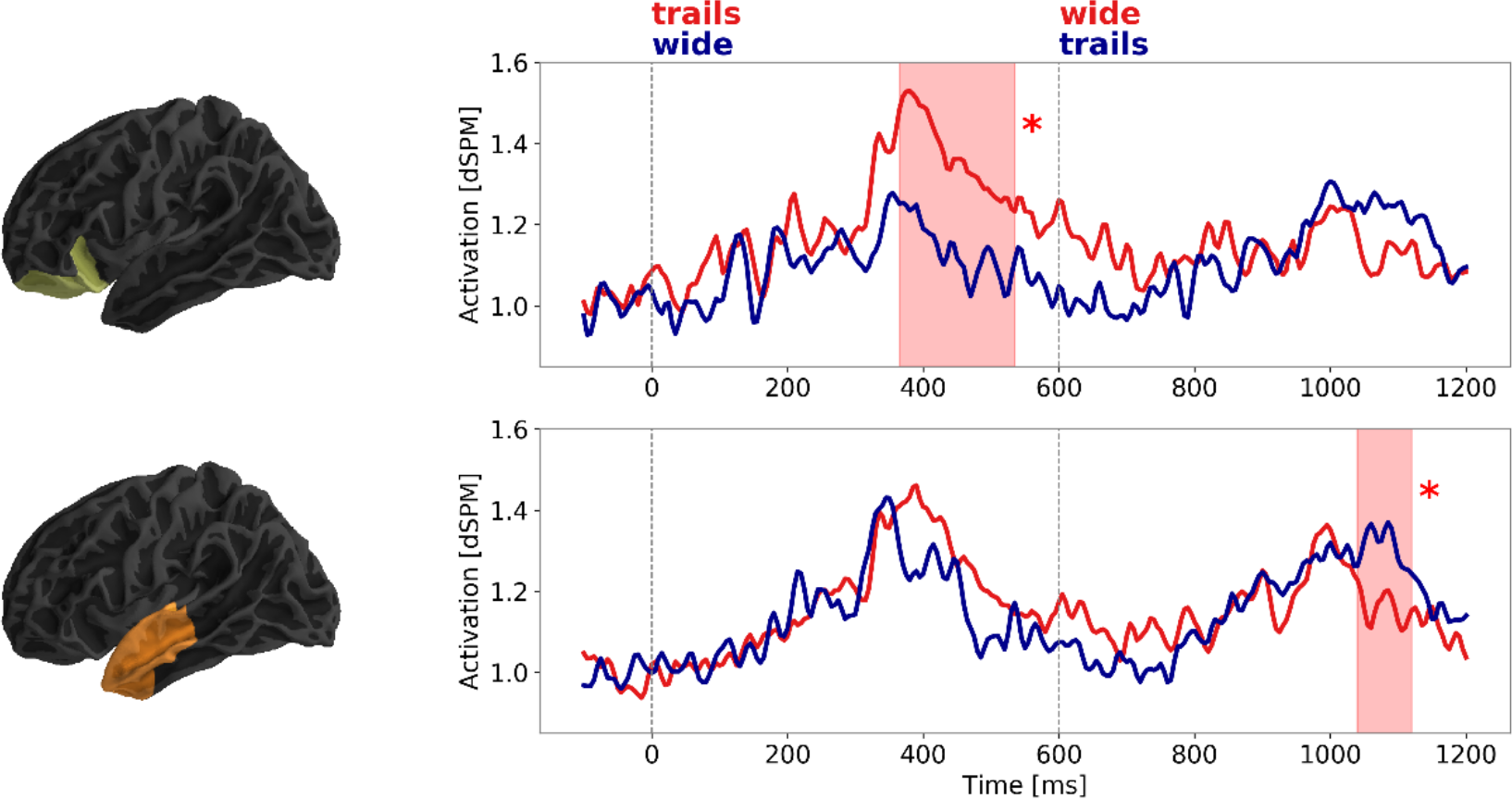
Repeated measures analyses of data from both experiments. Clusters capturing main effects of Word Order, indicating sensitivity to word class, were identified in the orbitofrontal and left anterior temporal lobe ROIs, where they appeared on the first and second word, respectively. In both cases, responses were greater during recognition of plural nouns than adjectives.

**Figure 6.**
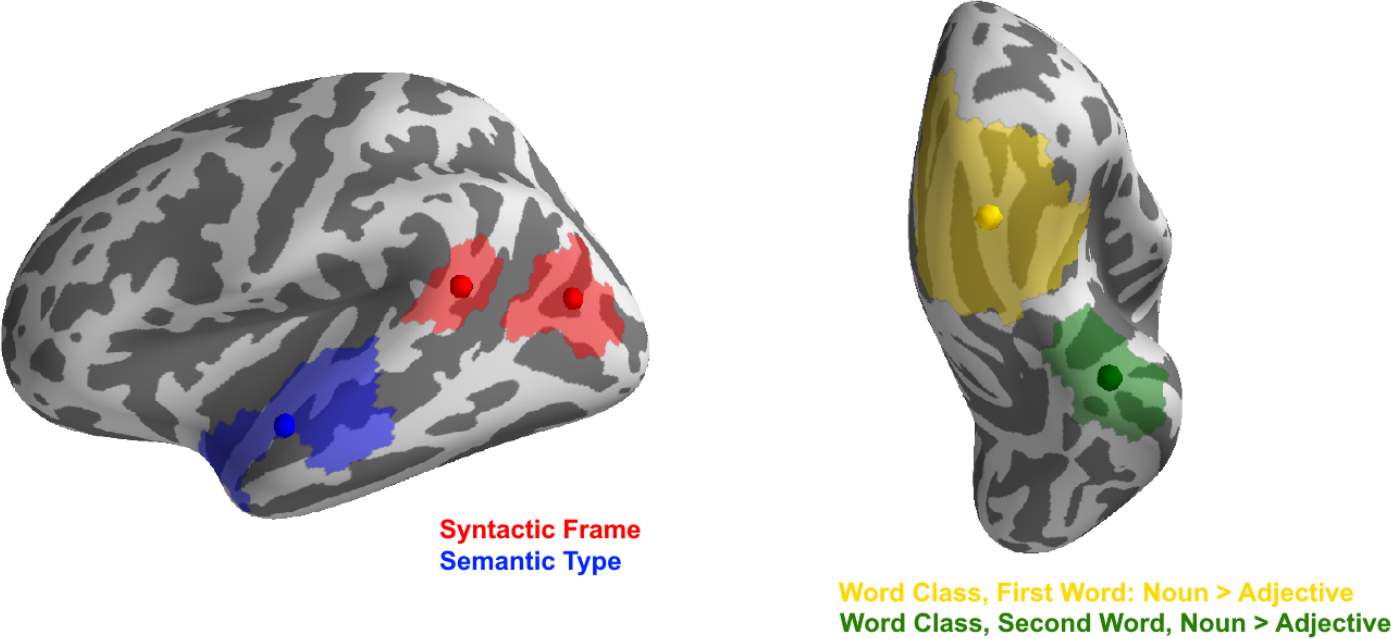
Spatiotemporal characterizations of the effects identified in the independent analysis of the Main Experiment (left) and the combined analysis of both experiments (right), displayed on inflated cortical surfaces. Patches indicate the spatial extent of clusters identified within the ROIs matching the effect in the original analyses. Spheres indicate the weighted center of this effect, based on mean F-values through time. In MNI coordinates, these centers were found at: Syntactic Frame (red): [-60, -48, 14] and [-41, -70, 13]; Semantic Type (blue): [-54, -6, -12]; Word Class, First Word (gold): [-39, 15, -35]; Word Class, Second Word (green): [-16, 35, -22].

Lastly, the combined analysis also identified a complex interaction in the orbitofrontal ROI in response to the second of the two critical words. This cluster spanned 400-445 ms post word-onset (p = 0.0176). In the Control Experiment, containing the adjective-noun word order, the cluster captured increased responses to event nouns in interrogative sentences, and entity nouns in declarative sentences. In the Main Experiment, the reverse was true: increases in response to entity adjectives in interrogative sentences, and event adjectives in declarative sentences. Due to the complex nature of this interaction, and the lack of any clear hypothesis that can account for the pattern, we refrain from interpreting this effect. All remaining clusters did not reach the threshold for significance (corrected p > 0.05).

### 3.4 Connectivity Analysis

We conducted our primary assessment of Granger Causality (GC) in responses to adjectives in post-nominal predication, which elicited an increased response relative to post-nominal modification. Significant GC values were found for the presumed connections from the PTL to the ATL (F = 0.0118, p = 0.0022), and the PTL to the IFG (F = 0.0115, p = 0.0027). Additionally, the GC value corresponding to the connection from ORB to the IFG was also significant (F = 0.0132, p = 0.0008). These patterns are summarized in Figure 7. The parallel GC analysis of responses to adjectives in post-nominal modification showed a different pattern of significance. Here, GC values were identified for the connections from IFG to ORB (F = 0.149, p = 0.0003), occipital cortex to the PTL (F = 0.118, p = 0.0021), and PTL to ORB (F = 0.0104, p = 0.0056).

**Figure 7.**
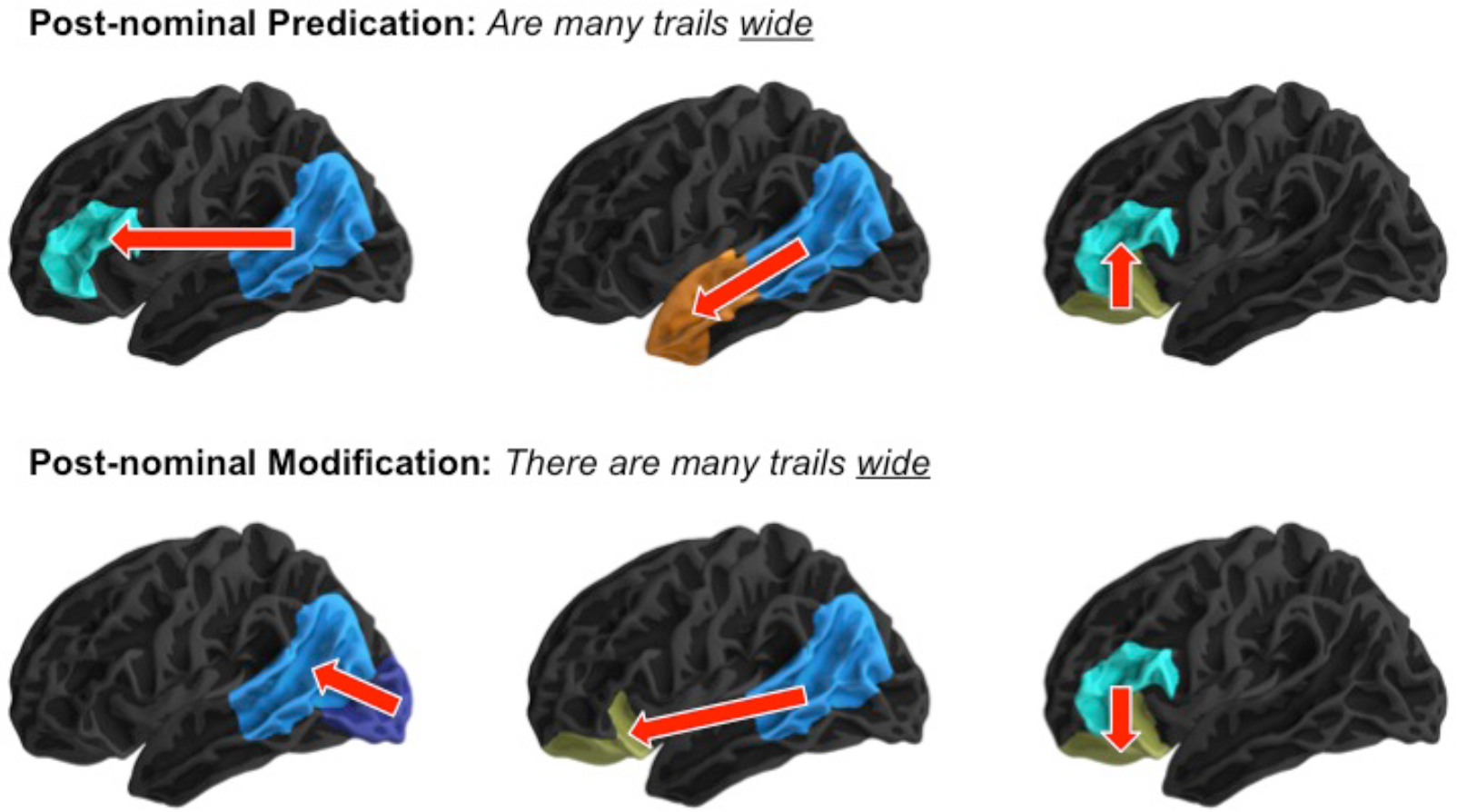
Granger Causality analysis of left hemisphere activity revealed three significant connections in the responses to adjectives (0-600 ms) in post-nominal predication: PTL to IFG, PTL to ATL, and ORB to IFG. In the parallel analysis of adjectives in post-nominal modification, significant connections were found from occipital cortex to PTL, PTL to ORB, and IFG to ORB.

## 4. DISCUSSION

Although a large body of research has attempted to elucidate the neurobiological basis of syntax, it has remained unclear whether purely structural properties of expressions can modulate neural activity in natural language processing. In other words, semantically unconfounded demonstrations of the presence of syntactic composition in the brain have been missing for normal, well-formed language. This gap in our knowledge has a principled cause in the fact that structure is difficult to vary without also manipulating meaning. Here we focused on a specific contrast of English syntax, post-nominal modification vs. predication, that allowed us to vary syntactic structure while keeping the lexical-semantic and conceptual properties of the expressions constant. Our analysis of responses to post-nominal adjectives in these two syntactic contexts revealed a modulation of activity in the left posterior temporal lobe, consistent with previous findings implicating this region in operations related to syntactic composition (Friederici, Rüschemeyer, Hahne, & Fiebach, 2003; Bornkessel et al., 2005; Pallier et al., 2011; Brennan et al., 2016; Matchin et al., 2017; Matchin et al., 2018; Zacarella, Schell, & Friederici, 2017; Rogalsky et al., 2018). The locus of this effect showed insensitivity to changes in semantic aspects of the composition, as would be expected of a syntactic component. The finding of increased PTL activation related to syntactic integration was accompanied by the results of a Granger Causality analysis suggesting a relay of signal, across a broader time window, from the PTL to the IFG and the ATL. The orthogonal manipulation of the semantic type (event vs. entity) of the preceding nouns modulated responses to post-nominal adjectives in the left anterior temporal lobe, at approximately the same time as the main effect of syntactic frame. These and other findings are discussed in further detail below.

### 4.1 Syntactic composition in the PTL

From prior literature, the primary candidates for basic syntactic combinatory operations have been the left posterior temporal and inferior frontal cortex, and both regions have often been found to activate together in response to syntactic demands. This comes from work that has manipulated syntactic complexity (Cooke et al., 2002; Bornkessel et al., 2005; Friederici, Makuuchi & Balhmann, 2009; Newman, Ikuta, & Burns, 2010), studied responses to syntactic violations (Friederici, Kotz, Scott, & Obleser, 2010; Friederici et al., 2003), compared instances of composition and non-combinatory controls (Vandenberghe, Nobre, & Price, 2002; Humphries, Love, Swinney, & Hickok, 2005; Zacarella, Schell, & Friederici, 2017), and used artificial stimuli (e.g., Jabberwocky) to isolate structural processing (Pallier et al., 2011; Matchin et al., 2017). Based on the pattern of these findings, particularly those focused on syntactic complexities or violations (see Friederici et al., 2003; Bornkessel et al., 2005) it has been proposed that posterior sections of the left superior temporal gyrus and/or sulcus are critically involved in processes related to the integration of syntactic and semantic information (see also Grodzinsky & Friederici, 2006; Den Ouden et al., 2012), while cortex in the left inferior frontal gyrus (particularly Brodmann’s Areas 44 and 45) underpin the basic construction of syntactic structure (Zacarella & Friederici, 2015; Zacarella et al., 2017; Friederici, Chomsky, Berwick, Moro & Bulhois, 2017).

In the present study, we manipulated syntactic composition within well-formed natural language expressions while controlling the possibilities for semantic composition, showing a sharp increase in left PTL responses to a post-nominal adjective in a context that allowed straightforward syntactic composition. This response was not modulated by semantic aspects of the composition. The use of MEG allowed us to characterize this activity both spatially and temporally, revealing a component that had a clear evoked nature at approximately 200 ms post word-onset. No increases related to syntactic composition were observed in the left inferior frontal gyrus, nor in the left anterior temporal lobe, matching the findings of a recent investigation of agrammatical comprehension that demonstrated atrophy to only the left posterior temporal lobe, not the ATL or IFG, could be reliably linked with the ability to process complex syntactic structure (Rogalsky et al., 2018). As such, the present results further suggest that the left posterior temporal lobe is a critical locus for the online construction of syntactic structure during language comprehension. This raises the question of why previous MEG and fMRI studies of phrasal composition (e.g., Bemis & Pylkkänen, 2011; Zaccarella & Friederici, 2015) have failed to consistently identify effects related to syntactic composition in the PTL. One possible explanation is that, contrasting that of the sentential material used here, the minimal structure of two-word phrases fails to tax syntactic machinery in a manner that produces the PTL’s response, and/or make it measurable via imaging and non-invasive electrophysiological methods. There is some recent evidence that is consistent with this possibility, with Matchin and colleagues (2017, 2018) demonstrating PTL increases in response to sentences relative to lists of words, but no such increases when comparing responses to phrases and word-lists.

While the present manipulation of syntactic composition perfectly correlated with a contrast in interrogative and declarative forms, a parallel experiment using near-identical lexical material in interrogative and declarative sentences failed to find any semblance of the effect that was observed for our manipulation of syntax in the main experiment (see Figure 3). This would thus appear to rule out an explanation of the increase as indexing a more general difference related to interrogative and declarative sentences. That said, as is always the case when the adoption of a conclusion hinges on a null effect, future work will be required to determine whether this pattern of results replicates.

Finally, the present results do not rule out the possibility that the left IFG also plays a crucial role in online syntactic composition. In the connectivity analysis of left hemisphere responses, we found an implied connection from the PTL to the IFG, alongside another to the ATL, that was present in the straightforward composition condition (predication) and absent in the other (modification), suggesting that the relay from PTL to anterior temporal and frontal lobe regions may be tied to the composition. While this rests on a qualitative comparison between the conditions, the hypothesis is compatible with previous evidence suggesting that a PTL to IFG link is critical to the processing of syntactic information, and that the loss of this connection leads to deficits in syntactic processing (Griffiths et al., 2013).

### 4.2 Semantic composition in the ATL

In the left anterior temporal lobe ROI, we identified an influence of the semantic type of the previous noun on responses to the post-nominal adjectives, 175-195 ms following noun onset. This finding that the left anterior temporal lobe is sensitive to semantic properties of the words entering composition is consistent with those positing a role for the ATL in conceptual representation and combination (Westerlund et al., 2014; Baron & Osherson, 2011). The relevant linking hypothesis is that a process akin to referencing this conceptual store takes place whenever lexical semantic properties are retrieved for composition. Here, we provide novel evidence that this characterization of the ATL holds in contexts beyond minimal composition designs (e.g., Westerlund et al., 2014), and fail to find evidence for a role of the ATL in online syntactic composition (c.f. Brennan et al., 2010).

Furthermore, the pattern of the semantic type effects identified here appears to be compatible with a more precise functional characterization of the ATL’s role in semantic composition. Using a minimal composition design (such as that described in Bemis & Pylkkänen, 2011), Ziegler and Pylkkänen (2016) demonstrated that the 200 ms ATL increase in response to phrasal composition was limited to instances when the meaning of the preceding adjective could be fully determined without requiring additional processing on the noun that followed. This led to the proposal that ATL activation at this time point indexes the building of ‘quick and easy’ conceptual representations. In the present results, in conditions where adjectives followed the nouns, the adjective-noun type pairing that elicited greatest amplitudes in the 175-195 ms window was the Entity-Entity match (e.g. *trails wide*). Thus, if one posits that this pairing, which represents the modification of physical properties of entities, is straightforward relative to the others, then the current pattern matches the quick and easy ATL characterization. Such a proposal requires an explanation as to why this pairing would be more readily composed at this point, particularly relative to event-event combinations. Another interesting question concerns how this developing characterization of the ATL’s role in composition relates to results showing correlations between this region’s responses to naturalistic stimuli and predictions from models of syntactic parsing (Brennan et al., 2010; Brennan et al., 2016; Brennan & Pylkkänen, 2017). We leave these questions open for future work.

### 4.3 Additional findings and limitations

The analysis of data from participants that took part in both experiments revealed two effects related to the order of the critical adjectives and nouns. Here, the left orbitofrontal cortex showed sustained increases in activation when the first of the two critical stimuli was a noun, relative to when it was an adjective. This effect spanned approximately 350 – 500 ms post word-onset. Likewise, on the second of the two words, the left anterior temporal lobe also showed increased activation in response to the nouns relative to the adjectives, 450-520 ms post onset. While several studies have focused on differential neural responses to nouns and verbs (see Vigliocco, Vinson, Drunks, Barber, & Cappa, 2011 for a review), we do not know of any work that has explicitly looked at differences in the processing of adjective and nouns. Furthermore, in their review of the literature, Vigliocco et al conclude that rather than grammatical class, the more important distinction in results from imaging and lesion studies is between object and action knowledge. Since it is unclear how this distinction may relate to the present findings, and there are many other explanations for differential responses to our adjective and noun stimuli (including the presence of a plural marker on the latter), we leave further consideration of these possibilities for future work.

Finally, we briefly point out two limitations of the present investigation. First, in our assessments of effective connectivity with Granger Causality, we used the entire time windows of responses to each individual word (i.e. 0 – 600 ms following word onset). This was chosen over the alternative of performing a multi-windowed analysis, or focusing on a more constrained time window, in order to use the maximum number of data points available. However, this does limit the conclusions regarding the critical points in the time series that gave rise to significant GC values. Future work may benefit from alternative means of assessing functional and effective connectivity that can not only identify critical windows for such connections, but also time-varying (i.e. non-stationary) relationships between regions. Second, although the visual presentation of the stimuli eliminated potentially confounding cues from prosodic structure and allowed us to determine the exact onset time of each individual word, an important question is whether the present findings relating syntactic and semantic composition to the PTL and ATL, respectively, hold when language is encountered acoustically. This thus presents itself as a straightforward means by which to validate the present findings.

## 5. Conclusions

The present investigation yields evidence from well-formed natural language expressions for a role of the left posterior temporal lobe (PTL) in incremental syntactic processing, while also providing a spatiotemporal characterization of activation related to this process. Cortex in the left PTL showed a robust activation increase approximately 200 ms following the onset of post-nominal adjectives, when these words could be composed with previous material to form a (temporarily) complete syntactic structure. Granger Causality analysis within a broader time window suggested that left PTL responses to these post-nominal adjectives were relayed to the ipsilateral IFG, which could support the notion that such a connection is critically involved in syntactic processing (Griffiths et al., 2013). Finally, we also found a separate influence of semantic type (related to the processing of entities and events) in the left anterior temporal lobe (ATL). The relevant results across two experiments can be accounted for by a model in which ATL activity at 150-200 ms post word-onset indexes the building of rough conceptual representations, working on semantic information that is quickly available following a word’s onset, while PTL activity at approximately 200 ms reflects a syntactic composition process. These results thus provide new insight into the neural mechanisms that support syntactic and semantic combinatory operations.

## Acknowledgements

This research was supported by grant G1001 from the NYUAD Institute, New York University Abu Dhabi (LP).

## Appendix A

**Table A.1.**
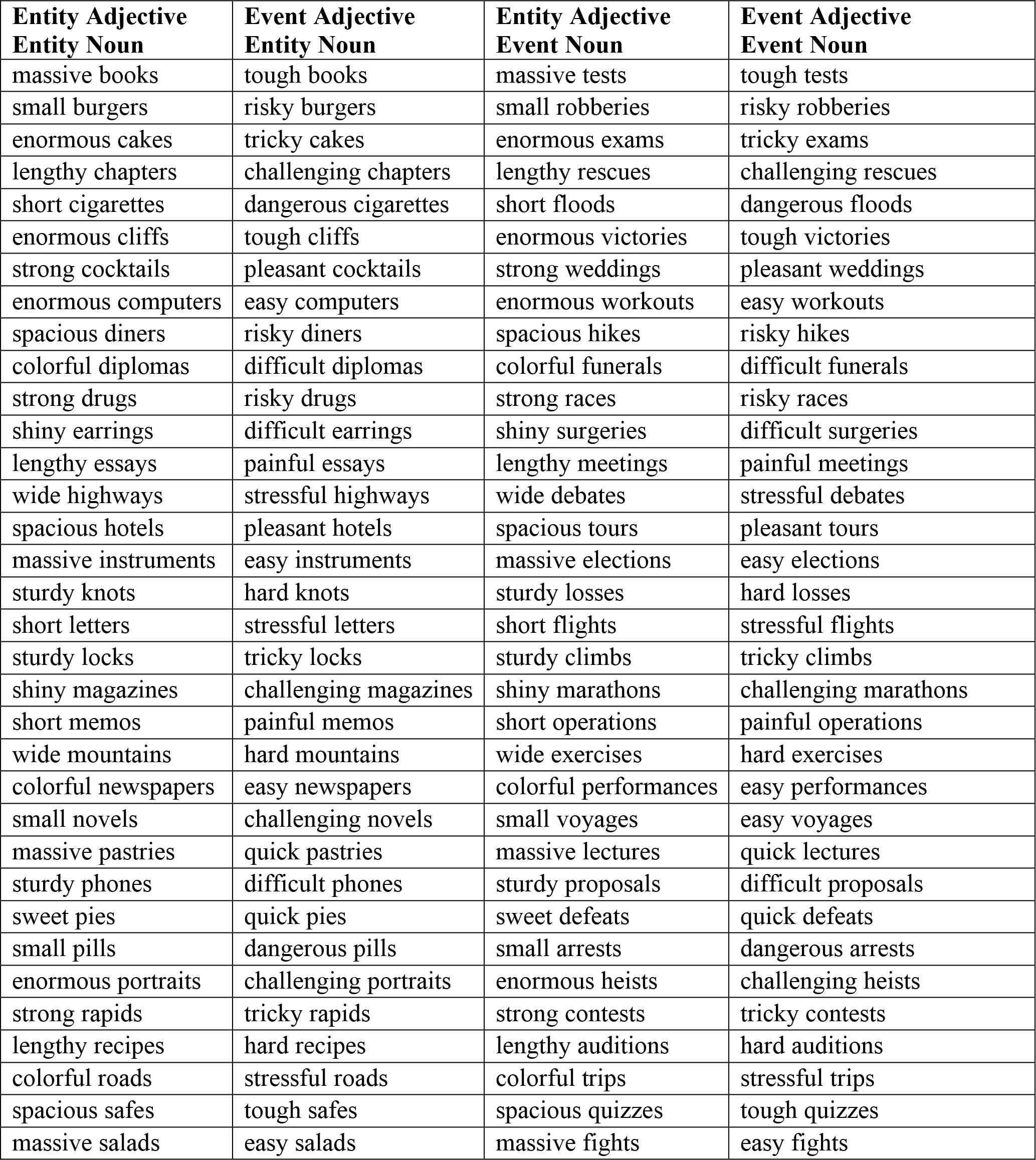
The complete set of adjective-noun pairs, organized in alphabetic order according to the entity-denoting nouns.

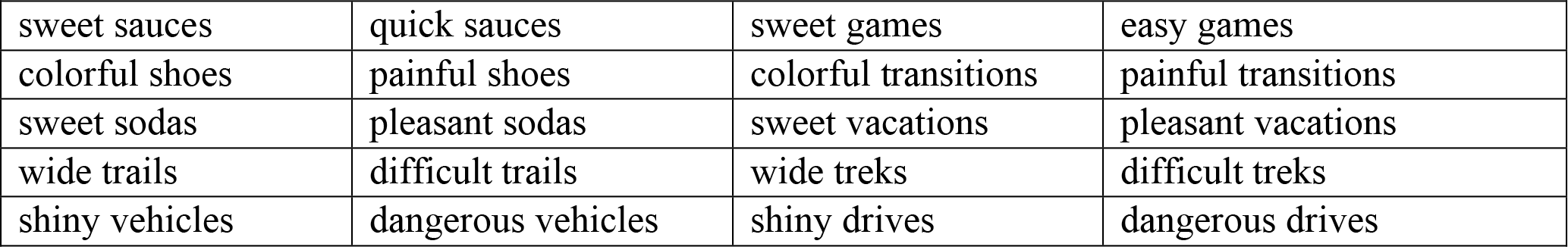

## Appendix B

We conducted two norming studies on Amazon Mechanical Turk (AMT) in order to confirm that our sets of nouns and adjectives did indeed differ on the desired dimensions. First, to confirm the entity and event denotations of our nouns, we embedded each noun in a sentence of the form “The [noun] happened last [month or season of the year]”. For example: “The trek happened last October“. Each AMT participant read a list of 78 sentences, containing each of the 78 unique nouns, randomly paired with one of the twelve months of the Gregorian calendar, or one of four seasons (fall, winter, spring, summer). Below each sentence, participants were presented with a 7-point Likert Scale, which they used to indicate how natural they found the sentence to be, with 7 indicating that they felt the sentence was completely natural, and 1 indicating that they felt the sentence was unnatural. Following rejection of participants who repeated the experiment, or failed to respond to all experimental items, each word received a rating from between 73 and 110 AMT participants (mean = 88.8). Consistent with the notion that they denote an occurrence in time, sentences containing event-denoting nouns as the subject of the verb “happen” were rated higher (mean rating = 5.77, standard deviation = 1.57) than sentences containing entity-denoting nouns in the same role (mean rating = 1.87, sd = 1.37). Inspection of mean ratings for each individual word revealed a clear dissociation between the two groups, with no entity-denoting noun receiving a mean rating above 3.04, and no event-denoting noun with a mean rating below 4.25.

The second AMT norming study was conducted to confirm that our event adjective, entity noun, pairings did indeed elicit an interpretation involving the insertion of an event, while the matching, entity adjective, entity noun pairs did not. To do so, nouns and adjectives were inserted into fill-in-the-blank sentences of the form:

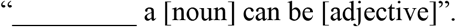

For example, in a coercive pairing, a participant would encounter the item: “[blank] a trail can be difficult”, while in a matching pairing, the item may be: “[blank] a trail can be wide”. Participants were instructed to fill in the blank in one of two ways: (i) If they felt that the material following the blank constituted a natural and complete sentence, they were to write the word “Complete” into the blank; (ii) if they felt that the sentence would sound more natural with a gerund-form verb in the blank (described as “an activity”, such as walking or hiking, with the possible addition of a preposition: “walking on”), they were to write in this activity. In this way, if the coercive pairs indeed elicited an event interpretation, they would cause participants to more frequently insert gerund-form verbs. Each participant was presented with a list of 13 unique adjective-noun combinations, such that no individual participant saw any word twice, with half coming from our event-entity condition, and half from our matching, entity-entity condition.

Following removal of participant repetitions, and trails on which participants failed to follow instructions, 19 to 30 unique participants responded to each word combination. An item score was calculated for each combination as the proportion of gerund insertions out of the total number of responses. Consistent with our design, mean item scores in the coercive pairings were higher, and near ceiling (mean score = 0.955, sd= 0.045), while matching pairs were lower, though also exhibiting greater variability (mean score = 0.325, sd = 0.161). Inspection of individual items revealed a robust tendency for coercive pairs to elicit an insertion, with the minimum item score at 0.842 (“tricky rapids”). Five matching pairs elicited scores greater than 0.50. In descending order, these were “sweet soda” (0.650), “short cigarette” (0.600), ‘colorful road’ (0.556), “sweet sauce” (0.533), and “lengthy recipe” (0.526). However, given the already small size of our stimulus set for an event-related MEG experiment (as regards the number of unique trials in each cell of the design), and the still clear dissociation of coercive pairs from matching pairs, we opted not to remove any items based on their scores. As such, following the norming results, this set of 78 nouns and 26 adjectives was confirmed as the final experiment material.

